# Biologically-informed Interpretable Deep Learning Framework for Phenotype Prediction and Gene Interaction Detection

**DOI:** 10.1101/2025.04.11.648325

**Authors:** Chloe Hequet, Oscar Gaggiotti, Silvia Parachini, Elena Bochukova, Juan Ye

## Abstract

The detection of epistatic effects has significant potential to enhance understanding of the genetic basis of complex traits, but statistical epistatic analysis methods are complex and labour intensive. In recent years, Deep Neural Networks (DNNs) have emerged as a powerful tool for modelling arbitrarily complex genetic interactions in relation to a phenotype; however, their utility is often limited by the challenge of interpreting their predictive reasoning. Although DNN interpretation methods exist, they are typically not designed for genomic applications, leading to hard-to-understand outputs with limited relevance to the field. To address this gap, we introduce GENEPAIR – a novel DNN interpretation framework designed specifically for genomic data, aimed at detecting putative associated gene-gene interactions for a phenotype of interest. Our approach offers several key advantages including model agnositicity, robustness to sample- and variant-level data variance, and flexibility to integrate varied domain knowledge into interpretable features. We demonstrate the efficacy of our method by applying it to a DNN trained on genetic variant data to predict Body Mass Index (BMI). The results of the analysis not only reveal single gene influences in close alignment with literature but also uncover previously unreported gene-gene interactions, demonstrating its significant potential for genomic discovery.

**Author summary:** Understanding how genes interact improves our understanding of how genetic pathways influence common diseases, potentially leading to new treatments. However, identifying these interactions is particularly challenging when a large number of genes are involved. Machine learning models, such as Deep Neural Networks (DNNs), excel at detecting complex patterns in data, but interpreting these patterns from trained networks remains a significant challenge. We have developed a novel framework to extract insights from a DNN trained to predict a trait, revealing how genes in the dataset may interact to influence the model’s predictions. Our approach is easy to incorporate or adapt to biological prior knowledge compared to existing methods, offering a powerful tool for discovering previously unknown gene-gene interactions.

## Introduction

One of the most significant challenges in modern biology and medicine is to understand the associations between the genetic variation observed across organisms within a species (their *genotype*) and the unique physical traits they exhibit (their *phenotype*). In particular, it is important to understand the non-linear interactions between genes, known as *statistical epistasis*. Mapping these interactions is fundamentally important for understanding the architecture of genetic pathways, quantifying genetic risk and highlighting potential treatment targets [1].

The most commonly used approach for analysing genotype-phenotype relationships is Genome Wide Association Studies (GWAS) which employ marginal association models to regress a phenotype of interest against the linear effect of one or more genetic variants [2]. Although GWAS have successfully identified numerous common, low-penetrance variants associated with complex traits [3], they are typically unable to detect non-linear effects. They are also challenged by the sheer number of genetic variants and strong correlations among them [2]. Secondary analysis of GWAS data set using tools such as Polygenic Risk Scores (PRS) [4] and Gene Set Enrichment Analysis (GSEA) [5] have further extended the clinical and research utility of disease-associated variants highlighted by GWAS [6]. However, these approaches also assume variants are uncorrelated and interact in a linear additive fashion [7]. Consequently, they fail to capture the intricate non-linear interactions and higher-order effects that are believed to contribute to a substantial fraction of the missing heritability in some traits [8–10].

Deep learning models, including DNNs, have been proposed as an alternative approach that can overcome some of GWAS limitations. They use representation learning to automatically identify complex patterns without requiring explicit parametrisation, making them well-suited for handling big, complex, high-dimensional data [11, 12]. The nonparametric nature of DNNs means they are able to model arbitrarily complex genetic interactions and do not need to rely on a linear additive model [13]. DNNs have been applied successfully to an increasing number of genomic tasks [14]. However, a major pitfall of DNNs across different domains is their lack of interpretability, rendering them as a “black box” [15]. Interpretation techniques including gradient-based (DeepLift, LRP [16]) and perturbation-based (SHAP [13, 17], LIME [18]) methods have been adopted to interpret deep learning models in genetic research. However, they have shown low agreement between identified influential variants and the existing domain knowledge [18], which limits their practical utility. Another way to ensure interpretability is to incorporate biological prior knowledge to design a network architecture. GenNet [19] is built on biologically plausible connections; that is, mapping single nucleotide polymorphisms (SNP) inputs to their associated genes in the hidden layer. However, this biologically informed network does not always outperform the phenotype prediction network with random SNP-gene associations [19]. This raises a critical question of how and when to use prior knowledge in phenotype prediction DL models.

In this paper, we propose GENEPAIR – a novel, biologically informed yet model-agnostic framework for interpreting deep learning models trained on genomic data. Our framework is designed to identify putative associated genes and systematically explore gene-gene interactions. The key novelty resides in leveraging the learning capability of DL without making strong assumptions and then integrating domain knowledge *only* at the interpretation phase to design biologically meaningful features for post-hoc interpretation. By doing so, a DL model is able to better capture interactions among genetic variants, which can then be identified at the later interpretation step.

To demonstrate the utility of our approach, we apply it to genetic data related to Body Mass Index (BMI), a highly polygenic phenotype with significant public health relevance [20]. BMI has been shown to have a heritability estimate of 40–70%, with numerous common and rare variants implicated in its regulation [21, 22]. Many BMI-associated variants map to non-coding regions, complicating efforts to translate these findings into actionable biological insights [23, 24]. Through our proposed framework, we identify BMI-associated genes and their interactions, shedding light on the regulatory mechanisms underlying BMI while addressing the challenge of nonlinear epistatic effects.

## Results

### Overview of proposed methods

Our interpretation framework is built on a machine learning model that has been trained to predict a phenotype of interest from SNPs; since the pipeline is model-agnostic, any model capable of capturing nonlinear interactions could be used here. The core innovation lies in the systematic feature ablation approach that probes the trained model to quantify the impact of specific perturbations on phenotype predictions. Typically, a method such as Gene Set Analysis may be used to map influential SNPs to genes and pathways, as these are more biologically informative than SNPs alone [25]. We similarly design interpretable features by mapping SNPs to their corresponding genes, referred to as *gene features*, resulting in DL network interpretation results that are immediately biologically useful without incorporating as many separate analysis steps. In the following, we introduce the design of gene features and ablation method and illustrate how we extend the method to explore gene-gene interactions. Details of the DL approach used in the first step of our framework are presented further below in the Materials and Methods section.

### Design of biologically-informed features

The goal of our study is to design a set of interpretable features that, when their interactions are investigated, provide more biological insight into the genetic architecture of the phenotypic trait under study than the original inputs, i.e., individual SNPs. Calculating attribution scores for each SNP suffers the same limitations as GWAS: namely, the effect sizes of individual SNPs in polygenic diseases tend to be very small, so this knowledge is not immediately useful for biological or clinical purposes. Additionally, considering only the interactions between lead SNPs ignores the possibility of higher-order interactions between multiple SNPs in multiple genes and again relies on further analysis steps to highlight potentially influential pathways. To generate immediately informative interpretation results, we use gProfiler SNPense [26] to group SNPs into gene features by linking them to genes based on location, a practice commonly used as part of variant annotation [27, 28]. While our approach focuses on genes as gene-gene interactions are typically of the most utility in terms of identifying functional pathways, the proposed method is flexible and inputs may be grouped into whatever features are of particular research interest. We do not perform any Linkage Disequilibrium (LD) pruning before either phenotype prediction or model interpretation as DL models are capable of adjusting input weights to account for variant correlation automatically. Much has already been discovered by GWAS and Gene Set Analysis about the genes associated with BMI [29], enabling us to use single gene ablation to validate the *reasoning* of the trained model against the established findings. With the knowledge that the network is accurately replicating biological knowledge, we may then move onto interaction detection.

### Feature ablation

Feature ablation is the process of perturbing a sample by replacing feature/s of interest with a baseline value and then quantifying the resulting change in model predictions [30]. In the context of gene features, this process is conceptually similar to a genetic knockout study, in which the expression of genes of interest in an organism is artificially reduced in a lab in order to see how each gene affects a trait of interest. In the synthetic case, SNPs are encoded using a 2D vector representing the number of reference (REF) and alternate (ALT) alleles; that is, [2,0], [1,1], or [0,2] for homozygous reference, heterozygous, and homozygous alternate respectively with GRCh37 (hg19) [31] as the reference panel. The effect of a gene feature is switched *off* by assigning all SNPs associated with the gene a baseline value of [0,0], corresponding to zero magnitude in both the reference and alternate allele vectors. If the resulting difference in output of the model due to a perturbation of a gene (referred to as the *attribution score*) is greater than expected, this indicates that the model uses this feature preferentially for prediction.

We begin with a model *g* trained on a training dataset 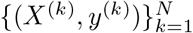, where *N* is the number of individuals and *X*^(*k*)^ ∈ ℝ^*D×*2^ is a SNP input with a dimension of *D. X*^(*k*)^ can be written as 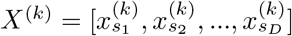,where each 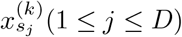is the [REF, ALT] encoded value for the *j*th SNP in the *k*th sample. *y*^(*k*)^ represents a value of a certain phenotype on the *k*th sample. We define gene feature set 𝔽= {*F*_1_, *F*_2_, …, *F*_*M*_}, where each gene feature *F*_*i*_(1 ≤ *i* ≤ *M*) corresponds to a set of input SNP features *S*_*i*_ ⊂ {*s*_1_, *s*_2_, …, *s*_*D*_}. Between gene features *F*_*i*_ and *F*_*j*_ there can be overlapping between their mapped SNP features; that is, *S*_*i*_ ∩ *S*_*j*_ ≠ ∅.

To learn about the attribution of gene features in 𝔽, we first obtain a prediction for a certain input *X*^(*k*)^ on the trained model *g* as follows:

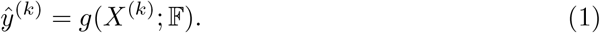

Next we evaluate the effect of gene feature *F*_*i*_ by setting the values of all its associated SNP *S*_*i*_ to their baseline and then recalculating the model’s prediction:

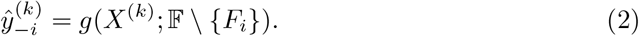

The *attribution* score of a gene feature *F*_*i*_ on the *k*th sample is the *difference* between the original and ablated predictions:

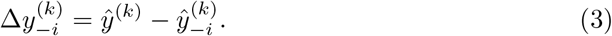

Feature ablation methods produce results at the individual sample level. To uncover aetiological trends it is necessary to aggregate results to produce an averaged attribution score for a sample population. The direction of effect for individuals would be different depending on the original genotype; for example, someone who is already homozygous for the risk alleles in *FTO* would see a decrease in BMI as a result of ablating the *FTO* gene, while the opposite would be the case for someone who does not have any of the risk alleles. To avoid confounding by the sign differences, we use the absolute values of the resulting differences for aggregation. The population-level attribution score of a gene feature *F*_*i*_ of a sample population *P* are thus aggregated by taking the mean absolute differences of all the samples in *P*; that is,

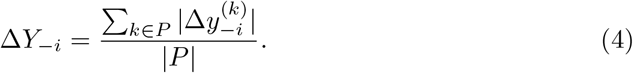

The attribution score indicates the effect of *F*_*i*_ in making predictions on a certain phenotype, providing a measure of the overall importance of *F*_*i*_ for the set of samples. To note that to test the generality of interpretation, the sample population *P* used for interpretation are often individuals in the test set that are not used for training the machine learning model.

Since our framework makes no simplifying assumptions about the relationship between *y* and *X*, it can detect the effects of any resulting relationship the original model has learned, regardless of complexity. However, this flexibility makes it difficult to determine a threshold for an *important* gene signal. To address this, we generate a null hypothesis distribution that assumes no relationships between a gene feature and the phenotype. Comparison with this distribution allows for quantifying the level of likelihood of obtaining a given attribution score if the null hypothesis were true (i.e., the p-value). To construct this distribution, we randomly permute the *y* values for the dataset in order to break any true associations between *X* and *y*. Then we train a machine learning model on this permuted data, which will produce ablation results representing random noise. On the off-chance that a particular network has learned a spurious correlation between one of the features and the permuted phenotypes, we have trained five random models and iteratively ablated the features to provide a truly random distribution of feature ablation scores. Since there are 1366 gene features in our study, 5 rounds of permutation and ablation result in a distribution of 6830 null attribution scores to compare against. Once a null distribution of gene feature effects is generated, this can be used as a cut-off value for real effects. Gene features producing effects that exceed the maximum value in the null distribution are considered significant, suggesting genuine relationships between the feature and the phenotype. This threshold ensure that identified signals are biologically meaningful and not artifacts of the model’s learning process. Figure 1 illustrates the ablation process.

**Fig 1.**
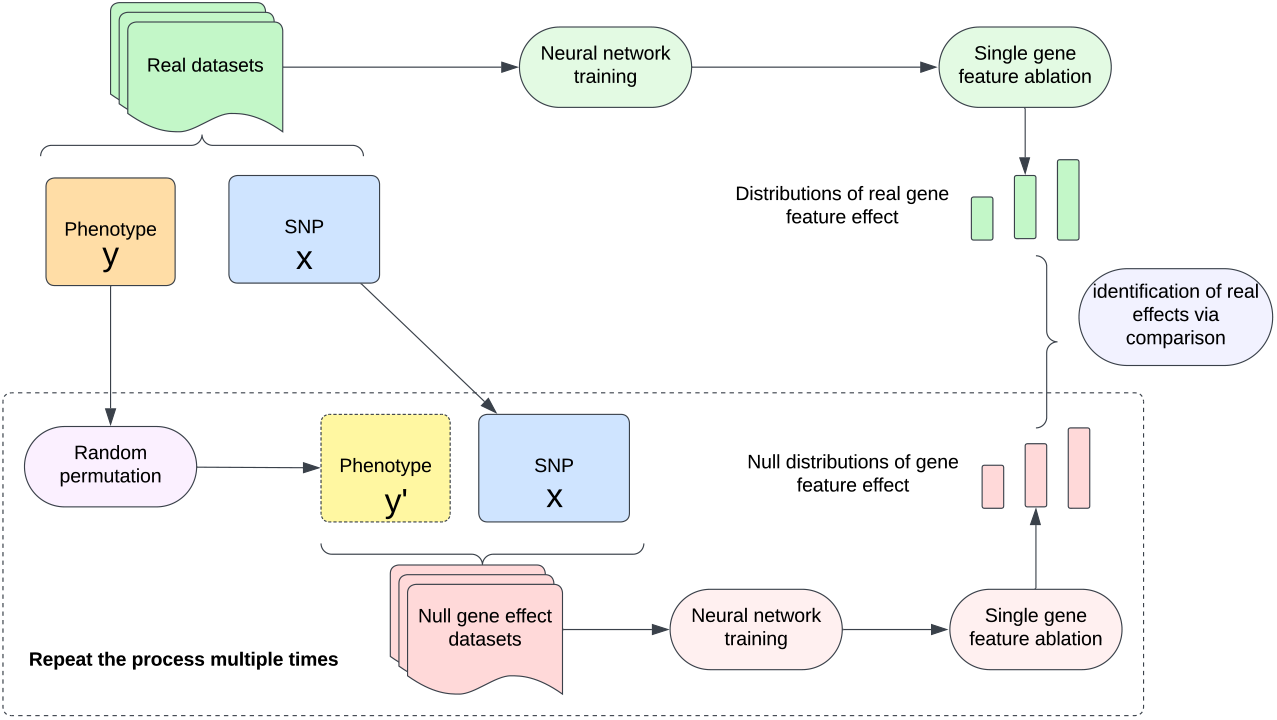
Illustration of the feature ablation process in GENEPAIR. The phenotypic contributions of individual genes are quantified by ablating each gene in turn and measuring the effect on the phenotype prediction by the trained network; this value is defined as the attribution score of the gene. A series of null phenotype predictors are created by performing random permutations the phenotype values and training identical networks on the permuted datasets. Per-gene ablation of these networks results in a null attribution score distribution against which the real attribution scores may be compared for significance.

### Gene pair feature ablation

Many individually important genes in BMI regulation have already been highlighted by GWAS and related statistical methods [29, 32]. Our single gene ablation method could hypothetically identify associated genes that are missed by GWAS if the constituent SNPs are individually uninfluential. Our next step is to explore potentially associated interactions, for which we utilise gene pair features to test for evidence of statistical epistasis. The simplest definition of pairwise interaction is any deviation from a linear additive model describing how two predictors *v*_1_ and *v*_2_ jointly predict a target variable *z* [33]. The simplest such linear additive model would be linear regression, in which a quantitative outcome *z* is modelled as a function of predictors *v*_1_ and *v*_2_ using the formula

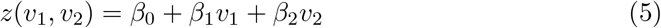

where *β*_0_ is a constant bias term and *β*_1_ and *β*_2_ are coefficients for predictors *v*_1_ and *v*_2_ respectively. Under this model, each predictor variable has a linear relationship with *z* such that a unit increase in *v*_*i*_(*i* = 1, 2) would result in a *β*_*i*_ increase in *z*.

To quantify deviations from the above additive assumption, a general model incorporating interaction, without making any parametric assumptions about the nature of interaction, is defined as

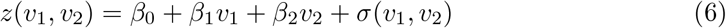

where *σ* represents an unknown interaction function, capturing any nonlinear, non-additive, or potentially higher-order relationship between *v*_1_ and *v*_2_. The null hypothesis *H*_0_ : *σ*(*v*_1_, *v*_2_) = 0 indicates that the relationship between the two variables and the output is linear and additive.

As with gene interaction, if there is no interaction between genes *F*_*i*_ and *F*_*j*_, we would expect that the contribution of the two genes together for a given sample *X*^(*k*)^ would equal the sum of the individual contributions, such that:

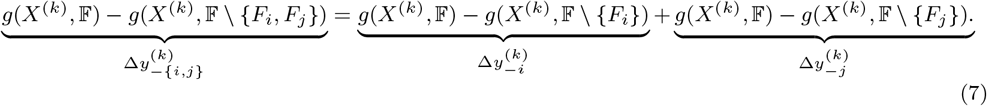

If interaction exists, the above equation no longer holds. The interaction term *σ*_*i,j*_ for a given sample *X*^(*k*)^ is defined as the deviation from additivity:

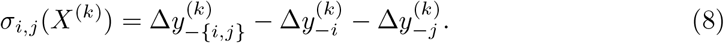

This can be conceived of as the effect on the value of *y*^(*k*)^ for sample *X*^(*k*)^ resulting from the difference in the individual and joint effects of *F*_*i*_ and *F*_*j*_, or the difference in *y*^(*k*)^ for sample *X*^(*k*)^ due to interaction between *F*_*i*_ and *F*_*j*_. No parametric assumptions are made about the nature of this interaction, but a nonzero difference in the individual and joint effects suggests an interaction. Zero of the value *σ*_*i,j*_(*X*^(*k*)^) corresponds to the truth of the null hypothesis, namely no interaction between features *F*_*i*_ and *F*_*j*_ for sample *X*^(*k*)^, but as with single gene feature ablation, generating a null distribution is necessary in order to reject the null hypothesis.

The procedure for generating a null distribution for interactions is similar to that for individual genes, except that now the null hypothesis is that there is no *interaction* between genes of interest *F*_*i*_ and *F*_*j*_, which may or may not be individually important. To obtain a null baseline it is necessary to interpret a model that has accurate main effects but no interaction effects; a well-performing linear model without multiplicative terms serves as an appropriate baseline for this purpose. By perturbing the linear model, a null dataset of interaction scores can be generated. This null distribution provides a baseline against which the observed interaction effects from the trained model can be compared. The interaction score for gene features *F*_*i*_ and *F*_*j*_ on a sample population *P* is the mean absolute score on all the samples in *P*;

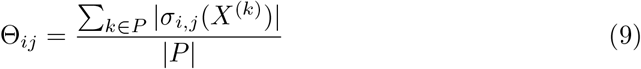

Significant nonzero interaction effects, as determined by comparing against the null distribution, suggest that the trained model has captured a genuine nonlinear relationship between *F*_*i*_ and *F*_*j*_. Figure 2 illustrates the pairwise gene feature ablation process.

**Fig 2.**
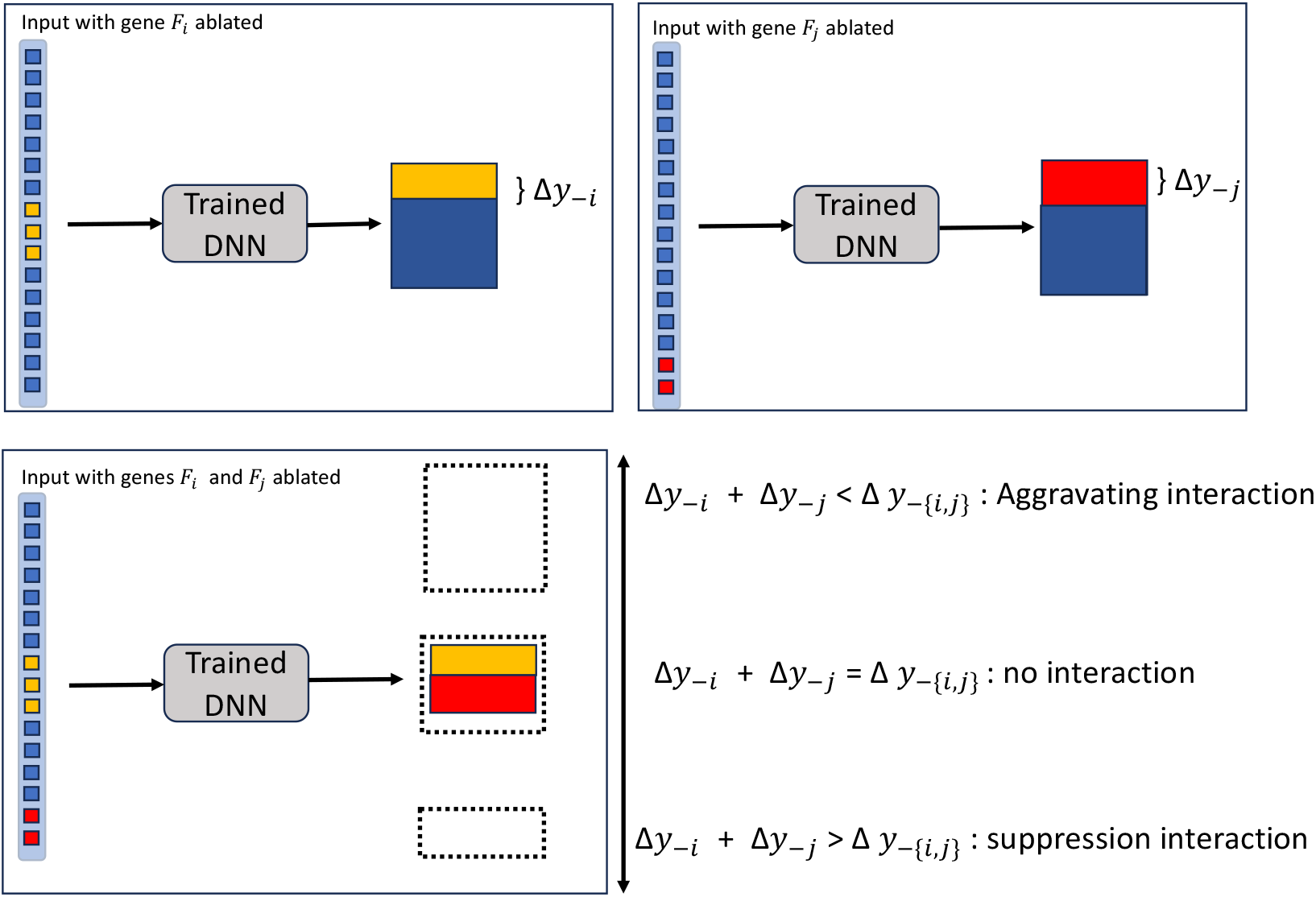
Pairwise gene feature ablation. Ablating each feature separately quantifies the phenotype attribution for the individual features. In the null hypothesis case of no interaction, the two effects combine in a linear additive fashion. A pair ablation attribution score that is greater or smaller than their linear effect suggests an interaction effect.

### BMI prediction

Our objective in building a deep neural network (DNN) for phenotype prediction is to avoid making strong assumptions while maximising the inclusion of SNP inputs within memory constraints. To this end, we perform GWAS on BMI in order to select input sets containing 10000 and 50000 BMI-associated SNPs, and assess the prediction performance using MSE (mean squared error) and Pearson R correlation coefficient (a measure of association between continuous variables). These results are compared against commonly used linear models, including Lasso and Ridge regression.

Figure 3 compare MSE and R scores between deep learning and the best performing linear model, which is a Lasso regressor trained with the Huber loss function. As the number of SNPs increases, the differences in performance between the linear baselines and the DNN become more pronounced, particularly in the 5,000 and 10,000 SNP sets. This trend suggests that larger SNP sets may increase the likelihood of detecting complex nonlinear interactions, which the DNN can effectively capture but linear models cannot. The DNNs have achieved the Pearson R scores of 0.247 and 0.271 with the 10,000 and 50,000 SNP input sets, respectively. These scores are close to the 0.292 Pearson R achieved using the PRS metric with 2.1 million variants [34].

**Fig 3.**
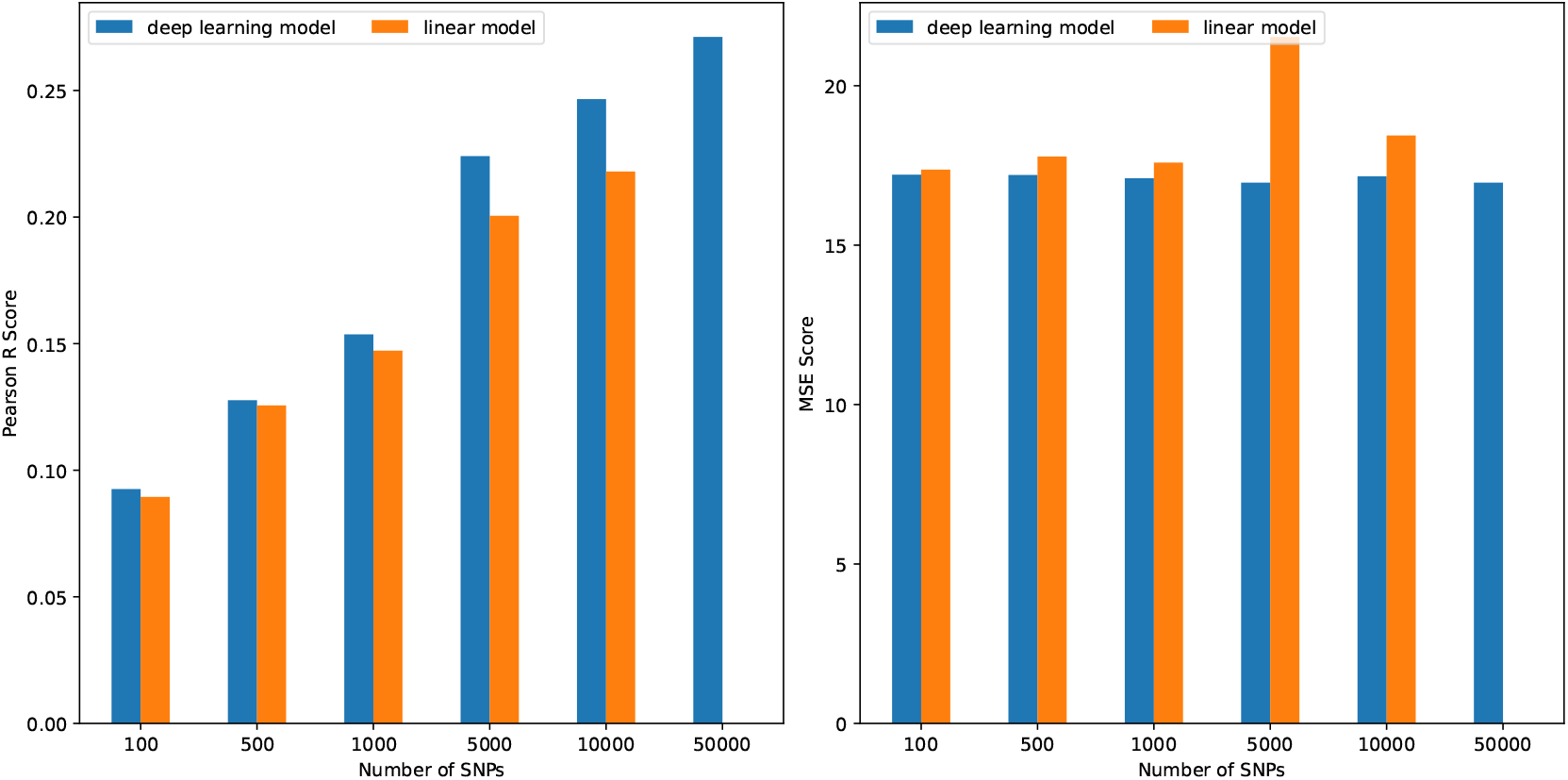
Comparison of MSE and Pearson R for the best performing linear and deep learning models on a different number of SNP inputs. Larger input sets improved Pearson R score for both models, but the effect was more pronounced in the deep learning model. The linear models were not able to utilise the 50,000 SNP input set due to memory constraints.

**Fig 4.**
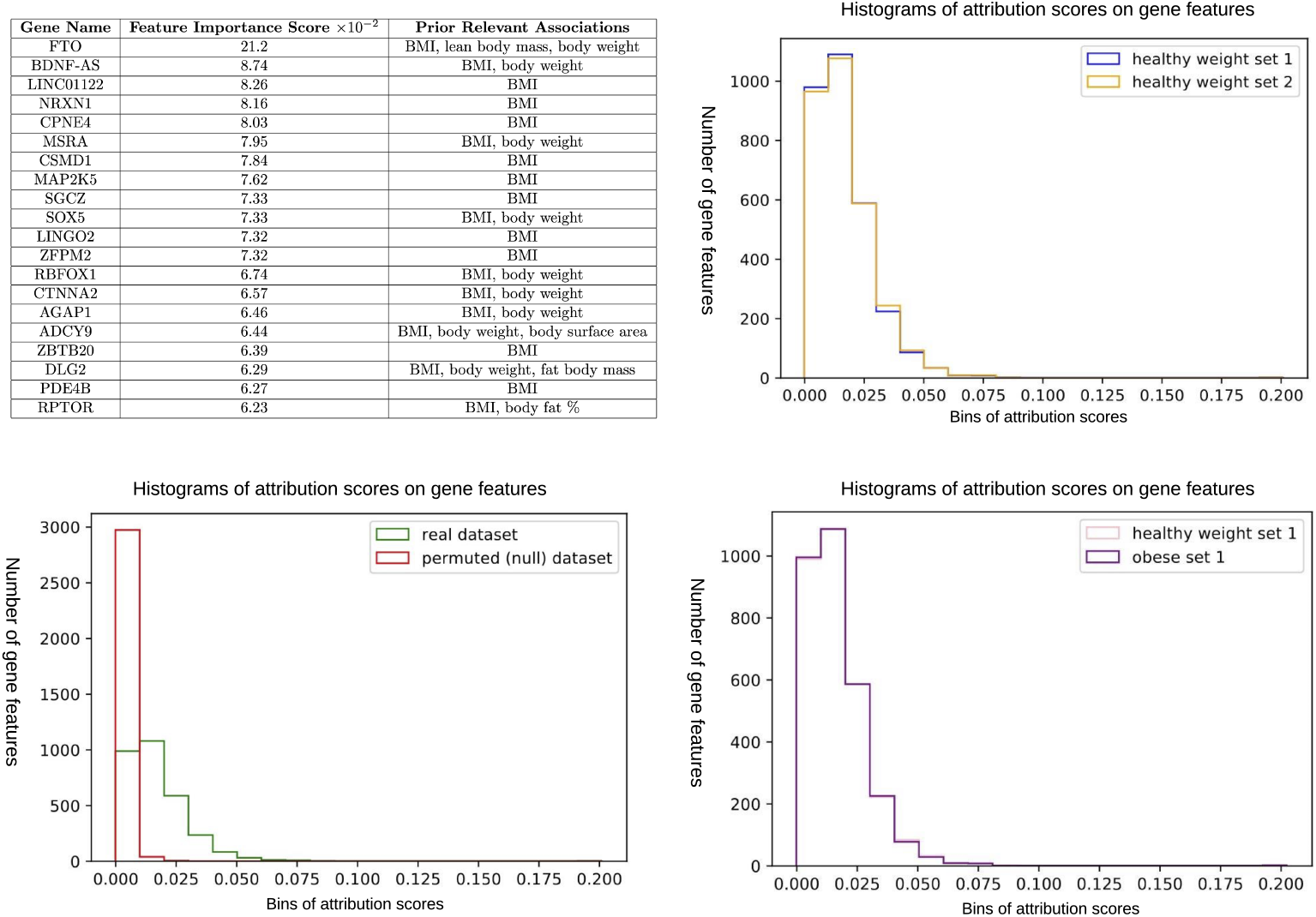
Single gene ablation results. All four test sets exhibit extremely close agreement in identifying the most important genes identified by single gene ablation. The top left table shows the genes identified in test set 1 using obese individuals; results from the other test sets are near identical with mild reordering. Prior relevant associations are sourced from the GeneCards database [36]. The two graphs on the right hand side show the attribution distributions of ablated genes on the two randomly partitioned test sets (top) and on samples from different BMI categories (bottom). Results from the other two test sets are omitted from these plots for legibility but can be found in the appendix. The close distribution match suggests a high level of agreement in ablation results between test sets 1 and 2, and between healthy weight and obese samples. The graph on the lower left compares the detected attribution scores in the real dataset with those detected in a network trained on the permuted dataset in which no genotype-phenotype relationship exists.

**Fig 5.**
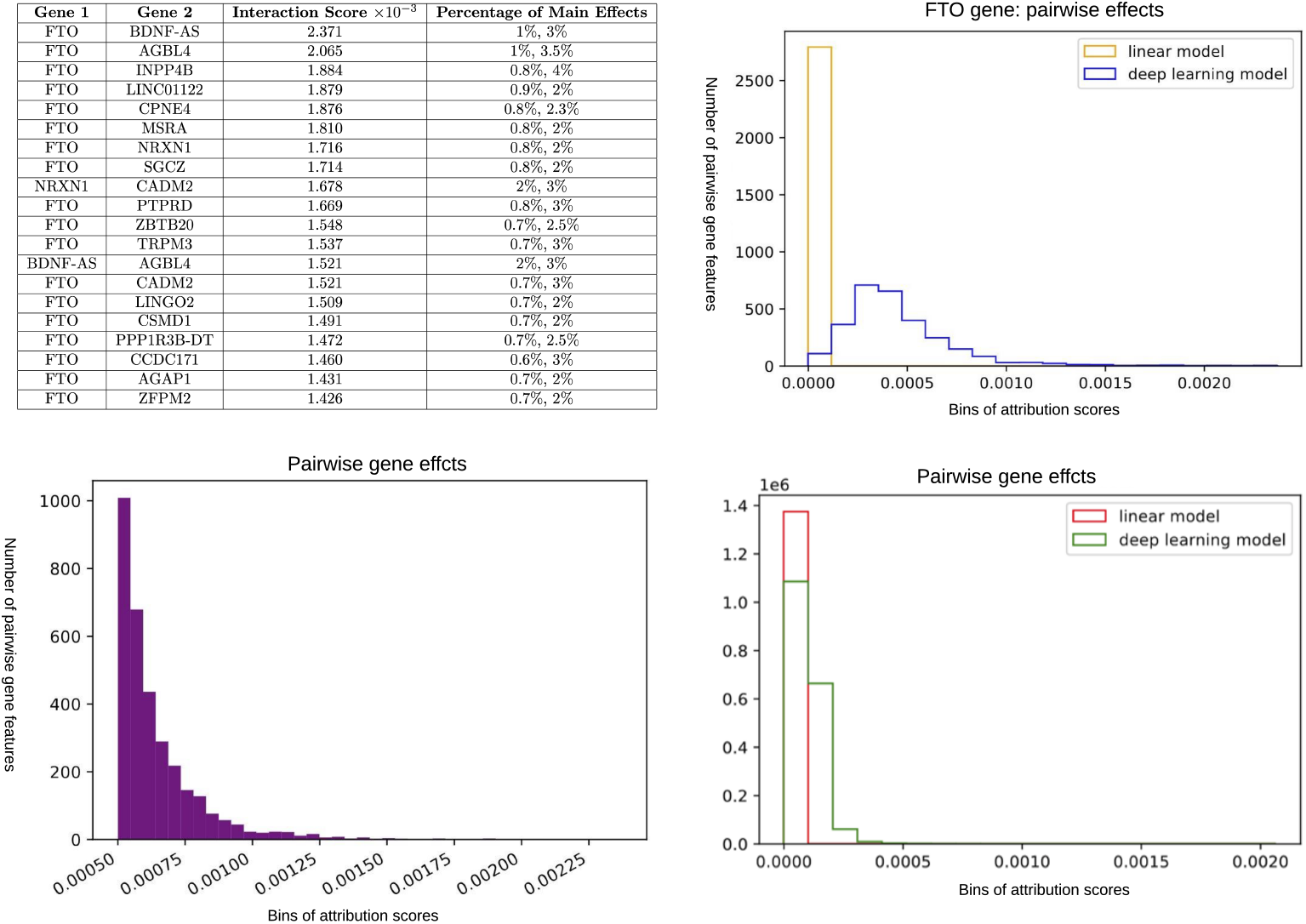
Results of pairwise gene ablation. The table shows the top pair effects detected in the MSE-filtered discovery set. The interaction difference corresponds to the average difference between the predicted BMI value difference that results from ablating both genes in the pair simultaneously and the result of adding the BMI differences of the individual genes. The bottom right graph compares the pair effects detected from the linear model with those detected from the deep learning model via gene feature ablation. The bottom left graph shows the distribution of pair effects greater than 0.0005. The top right graph shows the distribution of pair effects involving the FTO gene, as found by probing the linear model and the deep learning model.

**Fig 6.**
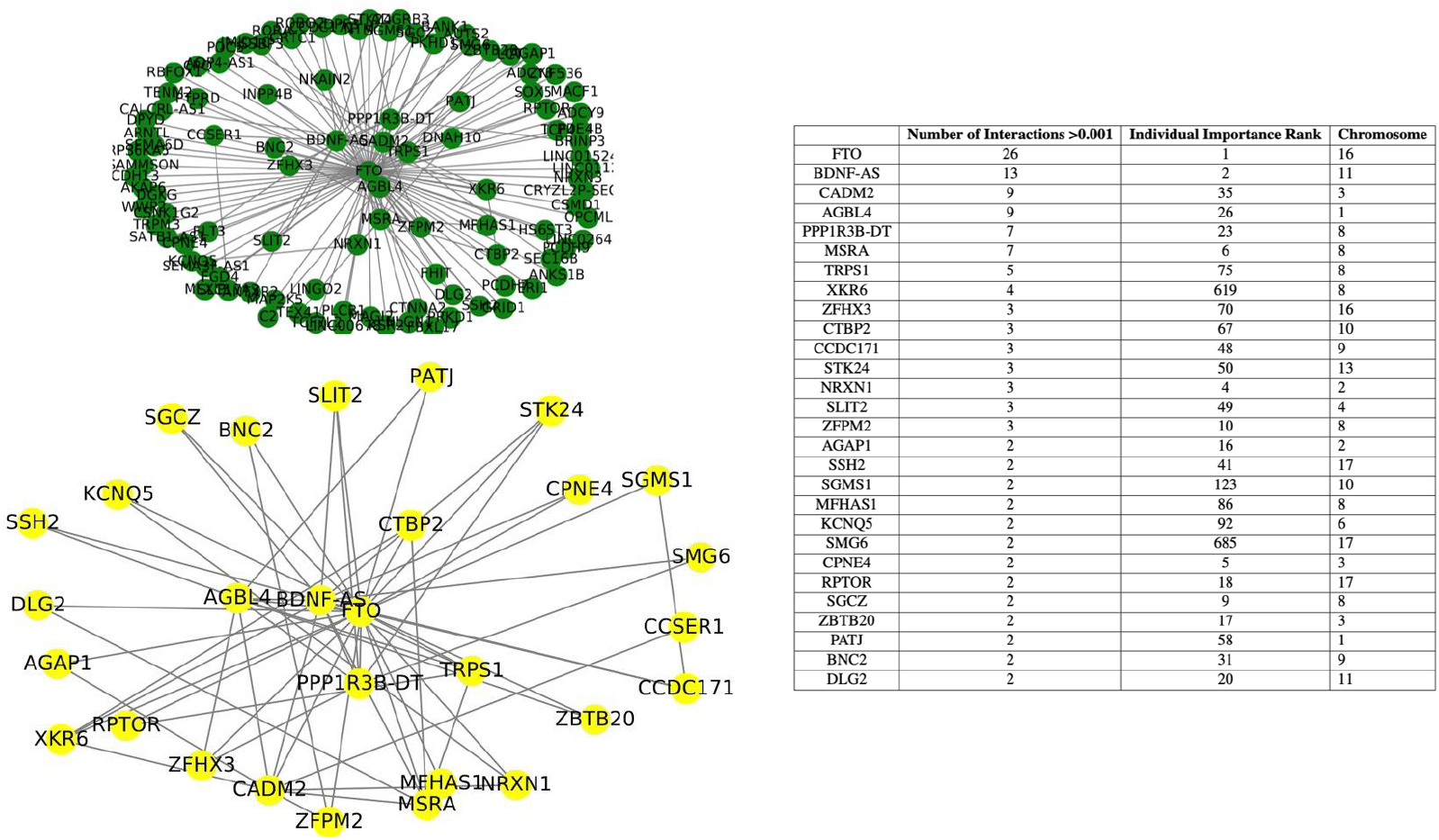
Gene interaction networks constructed with identified gene interactions. The table shows the top 20 genes with the highest pair effects greater than 10^*−*3^, in comparison with their individual importance rank. The top network illustrates all pair effects greater than 10^*−*3^, whereas the bottom network only shows genes with at least two such interactions.

### Associated gene identification

Our second objective is to identify phenotype-associated single gene features via the feature ablation method. This validates that the network has learned a good genetic representation of the phenotype and also identifies candidates for interaction. To assess the generalisation of our method in response to different populations, we stratify individuals in test samples by two BMI subcategories [35]; that is, underweight/healthy weight and overweight/obese. To ensure robust results, we only retain the samples for which the model achieves with a high level of prediction fidelity (prediction RMSE cutoff of 0.1) in an attempt to create a fairly homogeneous set. Statistics are then aggregated across each of these subsets to produce an average attribution score for each interpretable feature. We also randomly partition the healthy weight and obese datasets into two random subsets (test sets 1 and 2) respectively in order to explore the level of variation between individual samples. If two randomly selected subsets produce very different results, this suggests a high level of heterogeneity in important genes at the individual sample level.

In all sets, the *FTO* gene stands out with an attribution score that differs by a factor of ten, or greater than fifteen standard deviations, from the mean attribution score across all gene features. The attribution score of the *FTO* gene is equivalent, for a person of average height, to a 600g weight difference. This is consistent with the literature that the greatest weight difference caused by the FTO gene (the difference between homozygosity for the protective allele vs the risk allele) has been found to be about 3kg [37].

The second most important gene, BDNF-AS, is the antisense partner of BDNF, a brain-related gene shown to be heavily associated with severe monogenic obesity [38, 39]. The network identifies BDNF-AS as more important than BDNF, raising the question about whether the network has inferred information about BDNF indirectly through BDNF-AS, which has broader SNP coverage. Specifically, BDNF-AS is associated with 54 SNPs, while BDNF has 16 associated SNPs, of which all but two overlap with BDNF-AS. This highlights a potential limitation of using databases such as KEGG for grouping SNPs, as it may lead to certain genes *standing in* for overlapping counterparts. The remaining genes with high attribution scores have established associations with BMI and other related phenotypes, confirming that the network has, without deliberate input of such knowledge, automatically learned biological knowledge about BMI.

Analysis of the null distribution shows the vast majority of attribution scores are effectively zero, with the few higher attribution scores tailing off at 0.025. This suggests that any attribution score exceeding 0.025 would have a p-value close to zero, corresponding to a vanishingly small probability of occurring in the null hypothesis case. These attribution scores are likely to represent a genuine gene effect. In our real test datasets, 615 genes (20% of the total being tested) exhibit the mean attribution score to BMI prediction greater than this threshold.

Ablation results show a high level of agreement between randomly selected partitions within each BMI category, suggesting that different samples produce similar important genes and hence the method is robust to data variation and should be able to generalise to new data. The results are also similar regardless of whether the chosen samples are from obese individuals or normal/underweight individuals, suggesting that the trained model appears to look for the same patterns in all individuals regardless of BMI categories.

When probed for main effects using single gene ablation, the linear regression model has shown a similar attribution score distribution to the deep learning model, demonstrating that it has adequately learned to model most of the main effects. Like the deep learning model, it finds a standout main effect for the *FTO* gene and yields a very similar mean absolute BMI difference as a result of ablating this gene, at the attribution score of 0.198 (contrasting with 0.201 for the deep learning model). This suggests that it is a reasonable model of the main gene effects in absence of interaction.

As expected, none of the individual gene attribution scores, including *FTO*, are particularly large. This aligns with the understanding that complex traits like BMI are influenced by many genes of small effect. In our next section, we will look into interaction effects.

### Gene interaction results

Pair interactions are analysed on the same MSE-filtered sample set as single genes. For both the linear and deep learning models, the majority of attribution scores of gene pairs are small. However, in the linear model’s pairwise ablation results, a few gene pairs show relatively higher attribution scores; i.e., ~ 10^*−*2^. These correspond to genes sharing a number of overlapping SNPs in the dataset, including sense-antisense pairs, genes with separately labelled promoters, and genes with a “second degree overlap” where both genes share SNPs with a common third gene.

Overlapping variants in gene features will always cause a deviation from the linear model regardless of whether true gene interaction exists. As a result, the linear model identifies these overlaps as interaction signals. No nonzero interactions are detected by the linear model beyond these false positive overlap interactions that are the only signals present in the null distribution. This enables us to confidently attribute all nonzero interactions in the deep learning model (that do not share either a first or second degree SNP overlap) to the internal logic of the deep learning model, as opposed to being an artefact of the ablation method. The largest attribution scores for non-overlapping interactions in the linear model results is 10^*−*4^. Thus, we consider any interaction with an attribution score greater than this to be potentially meaningful, as it would again have a p-value of near zero and be effectively impossible under the null hypothesis. Interactions with attribution scores greater than 10^*−*3^ are regarded as strong evidence of genuine interactions captured by the model in the dataset.

An exhaustive search of all combinations of gene pairs, leading to 3.64 million pairs, is performed, with approximately 40% showing no detectable interaction. Among these, 140 pairs exhibit attribution scores greater than 10^*−*3^, which is an order of magnitude greater than the largest attribution score detected by the linear model. These significant interactions represent roughly 0.003% of all pairs tested, suggesting that pairwise gene interactions of this magnitude are a rarity for this phenotype in this cohort. In contrast, 95% of single gene attribution scores are no less than 10^*−*3^, indicating that individual gene effects are generally much larger than the detected pairwise interaction effects in this cohort.

In order to make sure that results are not skewed by the choice of samples, we make an additional benchmark set of the same size out of other samples in the test set that has not passed the MSE filter, and exhaustively run the interaction analysis for every pair of genes including *FTO* in the gene set on this group of samples. We find an extremely high level of agreement between different sample groups, showing that the method is robust across different data. This suggests that the model has learned a relatively good representation of the common variant basis of BMI for all the samples, and that the poor prediction performance on some samples could potentially be due to confounding environmental or rare gene effects that do not show up in the data but do influence the phenotype.

Most genes involved in the most strongly interacting pairs also have significant individual attribution scores, and perhaps unsurprisingly *FTO* is a main player. Pair effects tend to be only a small fraction of the main effect strength, and there are a lot of repeating genes in the top pairs, suggesting a possible network of interacting genes. Investigation of individual genes with numerous strong interactions can provide insights into the mechanism of genetic interactions that influence BMI.

*FTO* appears prolifically throughout the top pairs. Plotting its pair scores shows an unexpected distribution in which it appears to interact with the vast majority of genes in the dataset. 96% of other genes in the dataset have non-zero interactions with FTO and *FTO* shows interactions of greater than 0.5 * 10^*−*3^ with 30% of all genes. In fact, if we sum all of the pair effects associated with the *FTO* gene, it appears to indicate that it is possible that all of the *FTO* gene’s effect derives from its interactions with other genes, which is an interesting possibility since there is still some debate around the exact mechanism of *FTO* on BMI [23, 40–42]. This result raises the question as to whether *FTO* could act as a regulatory gene at the centre of a BMI gene network.

A potential gene interaction network is visualised as a graph, where nodes represent individual genes and edges signify interactions between them. Nodes are then ranked by connectivity; that is, the number of connections they share with other nodes. When only considering pairs with interaction scores above *>* 10^*−*3^, a distinct network structure appears: a central interaction network surrounded by peripheral genes, each of which connects to only one other gene. This indicates that the outer ring genes may not be intrinsic components of the regulatory network, but only acted on by it.

To isolate the possible regulatory network, we redraw the graph with a constraint that only includes nodes with two or more connections. The individual importance of genes, as determined by single gene ablation, is variable, and the implicated genes are distributed throughout the genome. This indicates a complex interplay where genes of varying individual significance are integrated into a broader network of interactions.

To the best of our knowledge, the majority of the interactions identified in our study have not been reported on before. A notable exception is an interaction network involving *FTO* and *ZBTB20* [43] via the *NRTK2* signalling pathway. A previous study [44] has investigated gene-gene interactions in BMI using multifactor dimensionality reduction. However, due to data removed in our data filtering and SNP-gene association steps, there is no overlap between their identified gene pairs and those represented in our dataset, preventing direct comparison of methods. However, our set does contain the gene *MAP2K5*, which is found in one of their detected interactions. We detect a strong interaction (*>*0.001) between *MAP2K5* and *FTO. FTO* is excluded from the MDR study, so this interaction would not have been detected. MAP2K5 also appears as one of the highly interactive genes in our interaction network. Overall, these results highlight a compelling combination of known genetic effects and previously undocumented pathways influencing BMI. The findings suggest that our proposed method can offer a promising new tool for uncovering genetic interactions within predictive models.

## Discussion

This paper presents a deep learning-based interpretation framework that combines predictive capabilities of neural networks and biologically informed features to uncover insights into genotype-phenotype relationships. Our method demonstrates robustness in discovering phenotype-associated genes and novel gene-gene interactions across diverse population sets, providing a flexible, nonparametric and model-agnostic tool applicable to a broad range of datasets and phenotypes. We leverage biologically informative interpretable features without the need to make strong assumptions, facilitating the discovery of novel insights into complex trait aetiology. In our experiments with BMI, despite the small size of detected interactions, the findings carry implications for BMI aetiology, showing that the method has promise for illuminating potential disease pathways.

In our study, GWAS is utilised to filter SNP inputs, which may have excluded variants with minor individual effects but significant interaction effects. This potentially limits the model’s predictive power and ability to detect interactions, as interactions do not necessarily only involve individually significant SNPs. Incorporating SNPs beyond those identified as GWAS-significant could improve model performance. To do so, it could be worth considering alternatives to GWAS for dimensionality reduction, such as newer ML-driven SNP prioritisation tools [45], or even unsupervised methods like autoencoders [46].

Our study follows the common practice of mapping SNPs to genes by location; for example, the Framingham Heart Study [47] is used as a basis for many GWAS protocols [28], or dbSNP [48] uses a 2.5kbp window to map SNPs to genes. However, this method is a relatively crude way of linking SNPs with genes that may contain a significant degree of error [49]. Lack of clarity around best practice for SNP-gene labelling is a major confounder for gene set analysis methods as well, and results in findings that are limited by the current limits of KEGG, which is far from complete [50]. Future studies could avoid this pitfall by using a more sophisticated SNP-to-gene mapping approaches that integrate multiple sources of genomic and functional data to refine the associations between genetic variants and their functional contexts, e.g. FUMA [51].

Our gene features are constructed using prior knowledge via gene annotations, however, it is not necessary to restrict ablation features to genes already associated with BMI, as is often done when constructing the features for biologically informed neural networks [19]. This allows our method greater potential of discovering novel genotype-phenotype associations, as well as lowering the risk of including incorrect information. Since the utilised priors (gene-SNP mappings) are phenotype agnostic, this approach is also not skewed in favour of better studied phenotypes.

This paper proposes one way of augmenting feature ablation for genetic data, but this method could be implemented in a number of different ways. As discussed, we used genes as interpretable features with the goal of pinpointing the most useful model explanation in terms of functional pathways, but the choice of grouping is up to individual preference. Our key contribution is a flexible pipeline allowing DL model interpretation based on the biological features of greatest potential interaction interest. With respect to baseline value configuration, it is not easy to infer effect direction from the proposed method due to the fact that when SNPs are ablated, the resulting direction of BMI difference will depend on what SNP is there in the first place. This also means that in pairwise gene ablation it is not possible to indicate what kind of interaction is being detected, only that there has been a deviation from the linear model. An alternative baseline could be the modal genotype, which would mean that the ablation results would indicate the effect direction of the minor allele, but this could significantly reduce the effect of perturbation and hence make results less useful if samples are heavily imbalanced in favour of the major allele.

Another option could be to keep the zero baseline but to consider alleles, instead of SNPs, as features, meaning that each allele would have its own ablation step. For example, instead of being set to [0,0], a heterozygous SNP could first be set to [1,0] and then [0,1]. In this way, the effects on the model prediction would be measured individually, helping separate the reference and alternate allele effects. This could show a more granular picture of effect and make it easier to pinpoint dominant interactions. However, this would become quite complex when one is trying to ablate entire genes or pathways, and is complicated by the lack of phasing information in most SNP datasets. Our proposed method is simple to implement and unaffected by SNP distribution while still capturing much of the available information.

A major pitfall of perturbation-based methods in general is that they are prone to problems when features are interdependent because impossible combinations can be generated by accident. Our proposed ablation method avoids this by perturbing fewer features at once, but there is still the possibility of perturbations that generate impossible allelic combinations that the model would by default not have been exposed to during training, leading to unexpected prediction results confounding interpretation. In future this could be circumvented by designing interpretable features in a way that takes linkage disequilibrium into account, for example using haplotype blocks as interpretable features instead of genes, or by analysing interpretable features for correlation in some other way and implementing requisite co-ablation of correlated features.

## Conclusion

The era of Genetic Big Data has provided unprecedented new opportunities for learning about the aetiology of complex phenotypes; however, being able to take advantage of this data for scientific and medical research requires the development of appropriate tools. We propose a framework for interpreting deep learning models using biologically informed features and probing the models for nonlinear feature interactions. We use this approach to probe our best performing deep neural network for its most influential genes in BMI prediction, and find that these genes are aligned with the existing findings about the aetiology of BMI. This result demonstrates that our feature ablation approach can be used to validate that a model has learned biologically accurate information. We show that ablation can also be used with combinations of features to search for interactions. This enabled us to explore gene interactions influencing BMI in data from the UK Biobank, and then use our most prominent detected gene pairs to construct a gene interaction map, validating existing interaction results and suggesting possible new interactions.

## Materials and methods

### Ethics Statement

This project was granted approval by the UK Biobank [52] under application ID 52480. The research has been conducted in accordance with UK Biobank Ethics and Governance Framework and under the policies of the supplied Material Transfer Agreement. Genetic data from the UK Biobank was used in accordance with the consent of the UK Biobank donors.

### Data acquisition and filtering

We acquire BMI data from the UK Biobank [53], a biometric database containing detailed health and genomic data for half a million UK-based adults aged between 40 and 69 years when recruited in 2006-2010. The Biobank holds extensive and detailed genotype and phenotypic information about its participants, with the goal of improving the understanding of the basis of common diseases, which are often complex and multimodal in nature.

Following the example of existing studies [54, 55] we apply sample quality control measures to exclude individuals of non-European ancestry and individuals whose reported sex does not match their biological sex from further analysis. Ancestry is determined by self-report of Biobank participants. We follow the UK Biobank’s exclusion recommendations to exclude samples with unusually high heterozygosity, samples with unusually high genetic relatedness to other samples (which could also include duplicate samples), and samples exhibiting sex chromosome aneuploidy. We also exclude individuals who are not used in the UK Biobank’s PCA calculations, and with NaN BMI values or a call rate of less than 95% [56]. Our final dataset contains 310471 distantly-related individuals of European ancestry. We randomly split the individuals into 80% for training (i.e., 248376) and the remaining 20% for testing (i.e., 62095).

### Genotype data preprocessing

SNP data in the UK Biobank was obtained using a custom Affymetrix Axiom array which directly measured 850,000 variants. From this, imputation was performed using the Haplotype Reference Consortium and UK10K + 1000 Genomes research panel, yielding in excess of 90 million SNPs however, due to memory constraints and concerns about DL model overfitting we only included directly genotpyed variants in our dataset and excluded imputed data. We conduct quality control by filtering out SNPs with Minor Allele Frequency (MAF) *<* 0.1% and missingness *>*3%, leading to 735221 SNPs.

SNPs are encoded using a 2D vector representing the number of reference and alternate alleles ([2,0], [1,1], or [0,2] for homozygous reference, heterozygous, and homozygous alternate, respectively). The reference panel is GRCh37 (hg19) [31]. Since it is computationally impossible to train a model on the entire SNP set, it is necessary to choose a selection of SNPs to train and test our models. A quantitative linear regression BMI GWAS is performed on the training set using Hail 0.2 [57]. Age, sex, and the first ten principal components were included as covariates. Full results from this GWAS can be found in the supplementary data. The most significant SNPs are used to create input datasets of 100, 500, 1000, 5000, 10000 and 50000 variants for the prediction models.

As shown in the Manhattan plot in Fig 7, the GWAS identified numerous significant peaks spanning the entire genome, but the most significant loci were located in chromosome 16. This is as to be expected in accordance with the literature [58] which identifies BMI as being highly polygenic but with the most significant association being found with the FTO gene on chromosome 16 [37].

**Fig 7.**
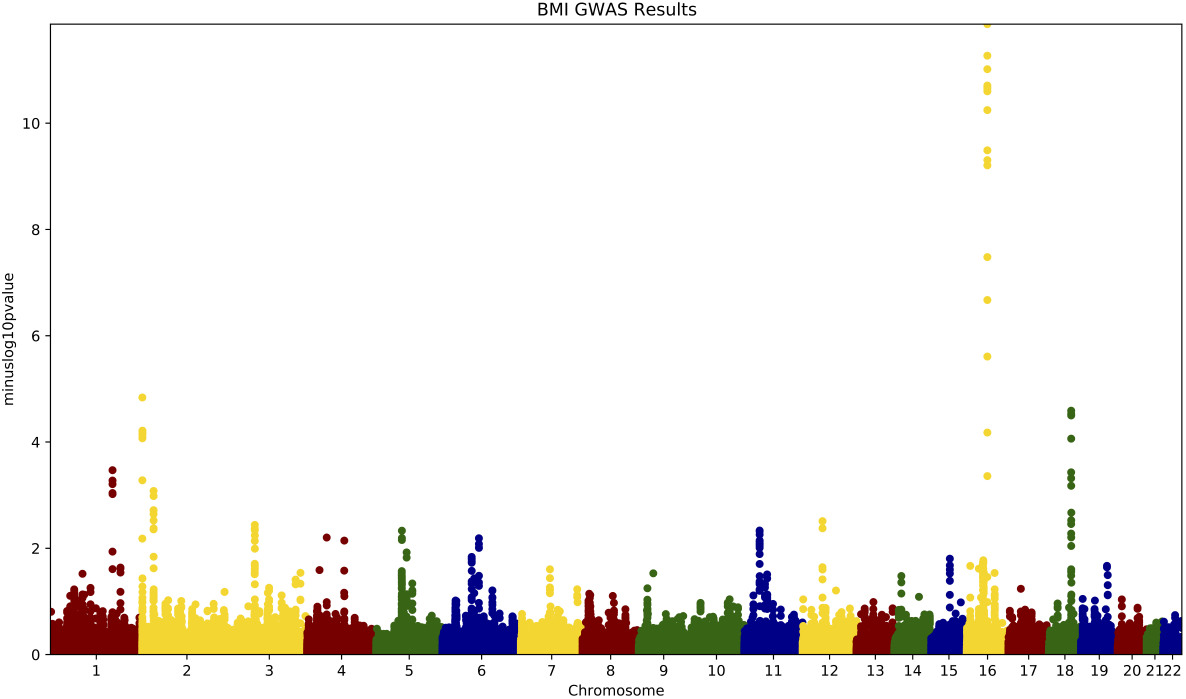
Manhattan plot showing the most significant SNPs discovered by our BMI GWAS. The y-axis shows the negative base ten logarithm of the p-value for SNPs in each chromosome.

### Linear baseline models

Due to the large size of the data, Stochastic Gradient Descent (SGD) was used to enable iterative training of linear models on the entire dataset [59]. A variety of different linear regression models were implemented by varying the regularisation penalty and loss function of the SGD Regressor. The loss functions trialled were squared loss, Huber loss, epsilon insensitive and squared epsilon insensitive, while the trialled penalties were L1, L2, and elasticnet. The best performing linear model was determined via gridsearch across all combinations of the above parameters. The best parameter combination was L1 and Huber loss on the 10,000 SNP set with a Pearson R score of 0.22.

### Neural network for phenotype prediction

As there is still no commonly agreed best neural network architecture for genotype-phenotype mapping, a large variety of different hyperparameters are trialled via grid search, performed with Ray Tune [60]. Following the state of the art methods [55], we use the Pearson R correlation between predictions and ground truth values on the test set. Both the Huber loss function [61] and Mean Squared Error [62] are trialled as loss functions, since MSE penalises outliers heavily and samples towards the ends of the BMI spectrum may be particularly genetically interesting. All Deep Learning models are constructed and trained with GPU-enabled Pytorch [63]. During training, the dataset was partitioned into 5 equal subsections, allowing the network to use each in turn as a validation set while training on the other 4 in a process known as cross-validation. By using the averaged scores across the 5 validation sets to determine network performance, we avoid the possibility of selecting a network that performs well on only a given train/validation partition but will not generalise well.

### Gene feature construction

Most SNPs are non-coding, so to group SNPs into related genes we employed the g:Profiler SNPense tool [64] and the KEGG [65] to link SNPs to genes using location and Sequence Ontology information, following the example set by many other studies [66–70]. SNPs not associated with any gene (i.e., 3101 in the top 10000 SNP set acquired from GWAS) are excluded from interaction analysis. SNPs associated with multiple genes (i.e., 937 in the top 10000 set) are included in multiple gene features, such that if either or both of the gene features were perturbed, this SNP would be as well. In order to be able to perturb samples it is necessary to have a way to switch genes *off* before feeding inputs into the original model, thus a baseline state of individual SNPs was encoded as [0,0], corresponding to 0 magnitude along either REF or ALT axes. This results in 1366 interpretable gene features instead of 10000 SNP features. To account for the possibility that different BMI subgroups may have different underlying genetic importance structures, samples are stratified by BMI category for the explanation phase, with samples with a BMI of 25 or greater being classified as overweight or obese in concordance with World Health Organisation guidelines [35]. These samples are selected from the holdout test set and have not been used at any point to train the model. The interpretation test set was randomly partitioned into two groups of equal size to aggregate results on separately in order to test the robustness of the method to changes in data.

## Code and Data Availability

Code to implement this method can be found on GitHub: https://github.com/chequet/genepair. Supplementary data, including full GWAS results, can be found here: https://zenodo.org/records/14919647. This research was conducted with data from the UK Biobank, which is available by application for *bona fide* research purposes.

## Acknowledgements

This work is part of the UK Biobank Project 52480.

